# Humanization of yeast genes with multiple human orthologs reveals principles of functional divergence between paralogs

**DOI:** 10.1101/668335

**Authors:** Jon M. Laurent, Riddhiman K. Garge, Ashley I. Teufel, Claus O. Wilke, Aashiq H. Kachroo, Edward M. Marcotte

## Abstract

Despite over a billion years of evolutionary divergence, several thousand human genes possess clearly identifiable orthologs in yeast, and many have undergone lineage-specific duplications in one or both lineages. The ortholog conjecture postulates that orthologous genes between species retain ancestral functions despite divergence over vast timescales, but duplicated genes will be free to diverge in function. However, the retention of ancestral functions among co-orthologs between species and within gene families has been difficult to test experimentally at scale. In order to investigate how ancestral functions are retained or lost post-duplication, we systematically replaced hundreds of essential yeast genes with their human orthologs from gene families that have undergone lineage-specific duplications, including those with single duplications (one yeast gene to two human genes, 1:2) or higher-order expansions (1:>2) in the human lineage. We observe a variable pattern of replaceability across different ortholog classes, with an obvious trend towards differential replaceability inside gene families, rarely observing replaceability by all members of a family. We quantify the ability of various properties of the orthologs to predict replaceability, showing that in the case of 1:2 orthologs, replaceability is predicted largely by the divergence and tissue-specific expression of the human co-orthologs, *i.e.* the human proteins that are less diverged from their yeast counterpart and more ubiquitously expressed across human tissues more often replace their single yeast ortholog. These trends were consistent with *in silico* simulations demonstrating that when only one ortholog is replaceable, it tends to be the least diverged of the pair. Replaceability of yeast genes having more than two human co-orthologs was marked by retention of orthologous interactions in functional or protein networks as well as by more ancestral subcellular localization. Overall, we performed >400 human gene replaceability assays revealing 56 new human-yeast complementation pairs, thus opening up avenues to further functionally characterize these human genes in a simplified organismal context.

## Introduction

Duplication of existing genes is regarded as a major contributing factor to the production of new genetic material [1]. Because of the immediate functional redundancy and dosage increase created following duplication, the most common fate of duplicated genes is loss of one functional copy, thus, returning to the ancestral state [2]. Rarely, however, duplicate gene copies either retain their ancestral roles or eventually diverge toward new functions. In the case of retained duplicates, several theories have been proposed to explain how multiple gene copies are maintained and their evolution post-duplication, though no clear consensus has been reached [3,4]. Sub- and neo-functionalization are two of the most commonly discussed theories regarding the evolutionary mechanisms leading to retention of duplicated genes. Generally, sub-functionalization posits that the function of the ancestral singleton is subdivided between its duplicates, leading to their mutual retention. Sub-functionalization can occur as a purely neutral process where degenerative mutations affect different subfunctions of both copies, resulting in their mutual retention in order to perform all functions of the ancestral singleton [5,6]. Sub-functionalization can also occur when duplicates acquire mutations that optimize subfunctions [7]. Neo-functionalization holds that the duplicated copy of a gene is freed from any strong selection, and becomes subject to drift until it is either lost through pseudogenization (a result sometimes termed ‘non-functionalization’) or acquires a new function that is fixed via positive selection [1].

Lineage-specific duplications also have a major impact on how homologous genes between species are related and identified. Two homologous genes related by gene duplication are termed *paralogs,* and are distinguished from those related by speciation, termed *orthologs* [8,9]. Importantly, an expanded family of paralogs in one lineage may be co-orthologs to a single gene in another lineage. How ancestral functions are partitioned, lost, or retained between paralogs and orthologs during these duplication events is a major topic of study for evolutionary biology. To address these questions, the *ortholog conjecture* was put forth, a major thesis stating that orthologs are more likely to retain function between species than paralogs, which will tend to diverge in function after duplication due to drift and relaxed selection [10,11]. This conjecture underlies common methods by which functions are assigned to newly discovered or understudied genes in un-annotated genomes [12], but tests of the conjecture have not been without criticism, at least in part due to the reliance on indirect functional assays [13–15].

In particular, direct experimental testing of the ortholog conjecture and retention of ancestral gene function is difficult, as neither detailed cellular and molecular function information nor consistent experimental assays are widely available for many genes across species, and much of the functional information assigned to new genes in understudied species is assigned by sequence similarity to genes of known function, further compounding the problem. Several studies have addressed these questions using comparative genomic or transcriptomic data as proxies for function, though they have been somewhat contradictory to date. One recent study in humans and mice found from GO annotations and microarray expression data that paralogs in the same species seem to retain similar mRNA expression patterns better than orthologs across species [13]. In contrast, an analysis of mRNA expression across several different human and mouse tissues suggested that orthologs retained function much more than their corresponding paralogs [16]. In an effort to reconcile these observations, Rogozin *et al.* compared expression profiles between human and mouse orthologs and paralogs and found that the apparent expression similarity of paralogs is conflated with species-specific background or batch effects, and that expression profiles of orthologs were significantly more correlated than paralogs after correction, providing support for the original ortholog conjecture [17].

A common limitation of such studies to date is that they use indirect proxy functional data (e.g. expression profiles) rather than direct functional assays. However, recent systematic studies of functional replacement of yeast genes by their orthologs from humans [18–21] and bacteria [22] demonstrate the power of inter-species gene swaps to directly test functional divergence and identify properties that determine functional conservation across vast evolutionary distances. Here, we sought to investigate the functional divergence across gene families by directly swapping human genes belonging to extended families into yeast. We identified all essential yeast genes in yeast-human ortholog groups (orthogroups) that have undergone expansions in the human and/or yeast lineage and systematically replaced the yeast orthologs with each of their human co-orthologs, assaying functional replaceability by complementation of a lethal growth defect. We find that duplicated human genes tend to differentially replace their yeast ortholog, rarely observing broad replaceability across members of expanded human gene families. Further, we quantified the ability of several protein-, gene-, or ortholog-based properties to explain the differential ability of human co-orthologs to replace and further support our observations with *in silico* simulations of protein evolution post-duplication. Collectively, our results suggest that within paralogous human gene families, at least one gene generally tended to retain ancestral function to a degree sufficient to replace a billion-year diverged ortholog in a yeast cell.

## Results and Discussion

### Identifying and selecting orthologs in expanded orthogroups and ortholog replaceability assays

Previous work from our group demonstrated that roughly half (47%) of one-to-one (1:1) yeast-to-human orthologs were capable of replacing essential yeast genes across a panel of three complementation assays (**Figure 1A**) [18]. We had specifically restricted these earlier tests to orthologs for which no lineage-specific gene duplications were easily identified, to mitigate effects of functional redundancy between paralogs. In the present study, we now focus on these cases of ortholog groups that have undergone lineage-specific gene family amplifications (**Figure 1A**). We again restricted our test set to include only those groups where the yeast gene was annotated as essential for growth under standard laboratory conditions (**Figure 1A, Supplementary File 1**) [23,24] and tested each human co-ortholog for its ability to individually complement loss of its yeast ortholog.

**Figure 1.**
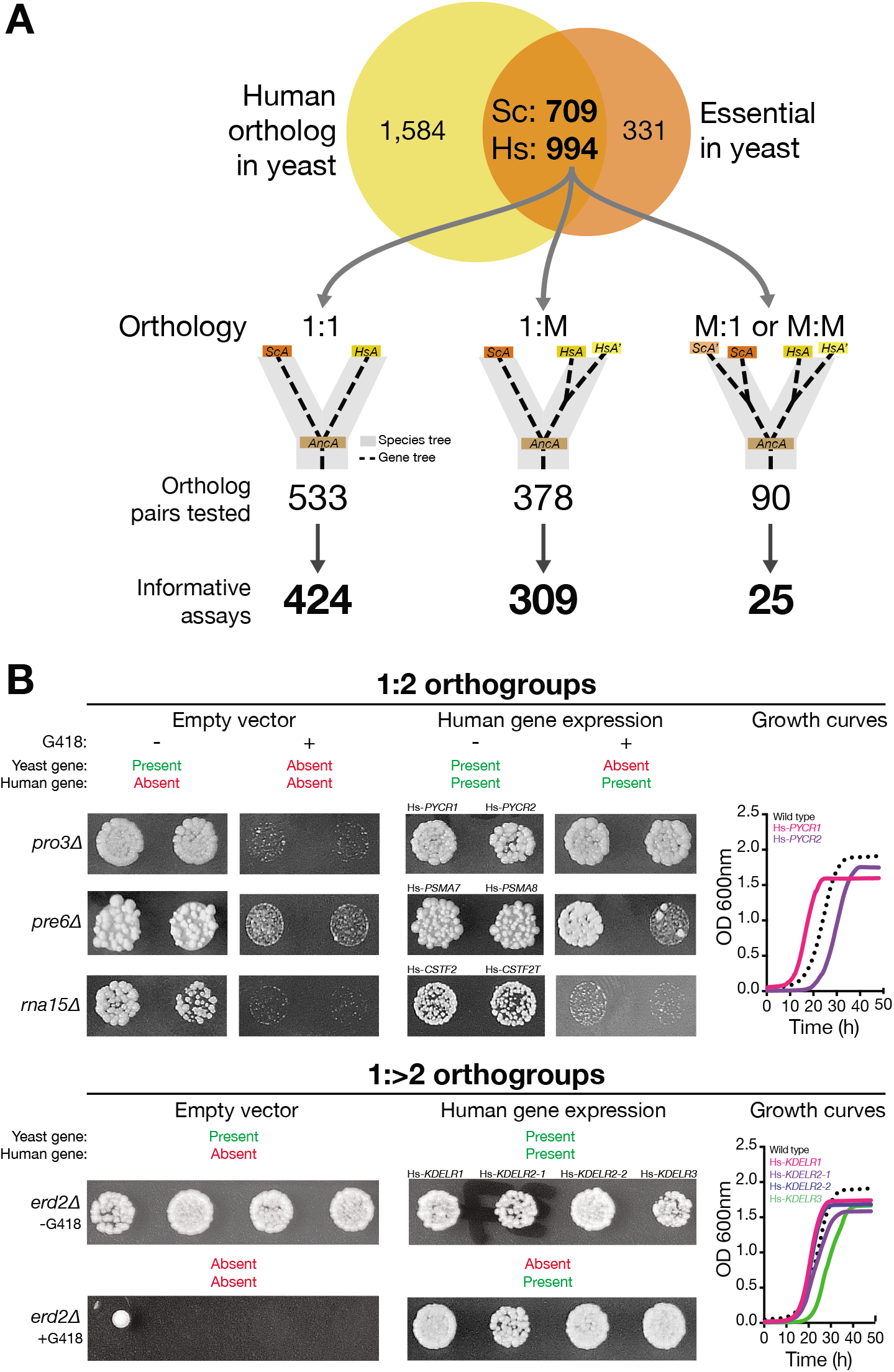
Systematic functional replacement of essential yeast genes with multiple human co-orthologs. **(A)** Of the 709 essential yeast genes with human orthologs, we had previously obtained results for 424 pairs with no duplications in either yeast or human lineage *(i.e.* with 1:1 orthology) [18]. In this study, we tested the remaining set of essential yeast genes that have acquired lineage specific duplications, classifying them as 1:M (1 yeast to 2 or more human co-orthologs) or M:M (>=2 yeast to >=2 human co-orthologs) or M:1 (>=2 yeast to 1 human ortholog). There are 140 essential yeast genes with more than one human ortholog, representing 378 ortholog pairs to be tested. In the case of 1:M category, we obtained 309 informative assays out of 378 testable pairs, whereas in the case of M:M or M:1 set we had 25 informative assays out of 90 testable pairs. Replaceability assays were performed in both heterozygous diploid magic marker collection and temperature-sensitive haploid yeast collection. **(B)** Representative assays performed in yeast heterozygous diploid magic marker collection for 1:2 (top) and 1:>2 (bottom) are shown. Magic marker heterozygous diploid deletion yeast strains expressing human genes were sporulated and the sporulation mix was spotted on magic marker media (see methods) with (yeast gene absent) or without (yeast gene present) G418. Assays were performed with empty vector control (human gene absent) or yeast expression vectors carrying a human cDNA (human gene expression). In the 1:2 class, three different outcomes of human gene replaceability in yeast were obtained: Top panel – both human co-orthologs (Hs-*PYCR1* and Hs-PYCR2) can replace their yeast equivalent (Sc-*PRO3).* Middle panel – one of the two human co-orthologs (Hs-*PSMA7* but not Hs-*PSMA8*) can replace their yeast equivalent (Sc-PRE6). Bottom panel – neither of the two human co-orthologs (Hs-*CSFT2* or Hs-*CSFT2T*) can replace their yeast equivalent (Sc-*RNA15*). In the 1:>2 class, an example of all human co-orthologs (Hs-*KDELR1*, two variants of Hs-*KDELR2*, and Hs-*KDELR3*) replacing their yeast gene counterpart equally well is shown. The yeast growth assays for these replaceable human genes is shown on the right for each. Haploid yeast gene deletion strains carrying plasmids expressing functionally replacing human genes (colored solid-lines) generally exhibit comparable growth rates to the wild type parental yeast strain BY4741 (black dotted-lines). Plotted growth curves display the mean of triplicate growth experiments.

In total, we identified 2,073 orthogroups using InParanoid [25], comprising 4,556 ortholog pairs (2,424 yeast proteins and 3,690 human proteins). Considering only yeast proteins essential for growth in standard laboratory conditions, there are 1,001 ortholog pairs available to test (**Figure 1A**) from 706 orthogroups. In our previous study, we obtained results for 424 human genes belonging to 1:1 orthogroups (**Figure 1A**) [18]. The remaining orthogroups comprise 468 additional pairs with essential yeast orthologs (**Figure 1A**) and are the focus of this study. These expanded protein families were further split according to having expansions only on the human lineage (one-to-many, 1:M) or those with expansions on either the yeast or both lineages (many-to-one or many-to-many, collectively referred to here as M:M). Based on the availability of appropriate yeast strains (see below and Methods), we identified 378 testable 1:M pairs, (involving 140 orthologous yeast genes), and 90 testable M:M pairs (36 yeast genes and 83 human genes) (**Figure 1A**). 1:M orthogroups varied widely in size, with the majority having only two human members, and the largest orthogroup (the melanoma antigen gene, or MAGE family) comprising 28 human co-orthologs (**Figure S1**).

We obtained human gene clones for our assays from either the human ORFeome collection or human sequence-verified Mammalian Gene Collection [26,27]. Human genes were subcloned *via* Gateway cloning [28] into yeast expression vectors under the control of the yeast GPD promoter, driving constitutive, robust expression. We performed complementation assays in one or both of two distinct yeast strain backgrounds: The first, referred to as the magic marker heterozygous diploid deletion (MM) collection, represents a collection of diploid yeast strains each of which contains a heterozygous diploid knockout of a single gene, with a selectable marker cassette allowing isolation of the haploid knockout post-sporulation [29]. The second mutant background is a library of temperature-sensitive (TS) haploid yeast strains, each harboring a mutant allele encoding a temperature sensitive form of the protein [30] that can be inactivated by growth at restrictive temperatures (typically 36 or 37°C). These two strain backgrounds allow precise conditional loss of an essential yeast gene or protein for assaying complementation by its human ortholog(s) (**Figure 1B, Methods**). We verified our assay results by isolating haploid humanized yeast gene knockout strains (either using magic marker medium or by tetrad dissection) while simultaneously verifying dependency on the human gene-encoding plasmid (**Figure S2, Supplementary File 1, Methods**). We further performed quantitative growth assays for each of the humanized yeast strains to more accurately characterize the robustness of complementation. The majority of complementing human genes exhibit similar growth profiles to the parental yeast strain (**Figures 1B, S3 and S4**).

In total, we obtained informative results for 309 of the 378 available assayable 1:M ortholog pairs, with results from at least one human ortholog for 131 of 140 (~94%) testable 1:M yeast genes. Of the 90 testable M:M pairs, 25 resulted in informative assays (**Figure 1A, Supplementary File 1**).

### Orthologs in expanded human gene families differentially replace their yeast orthologs

Notably, the rate of replaceability from the yeast gene perspective (*i.e.*, at least one human co-ortholog replaces the yeast gene) is similar to that previously observed for 1:1 orthologs (40% compared to 47%) [18]. From the perspective of tested human genes, 77 (~25%) of the 309 of the informatively assayed human genes could functionally replace their yeast ortholog, while 232 (~75%) could not (**Figures 2A, S3 and S4**). Where available, we compared our 1:M complementation results with published reports, observing a strong agreement at 92% precision and 96% recall (**Supplementary File 2**). In all, we report an additional 56 novel yeast-human complementation pairs. With previously published cases, this brings the total count of known human to yeast replaceable genes to 280 [18] (**Figure 2B, Supplementary File 1**).

**Figure 2.**
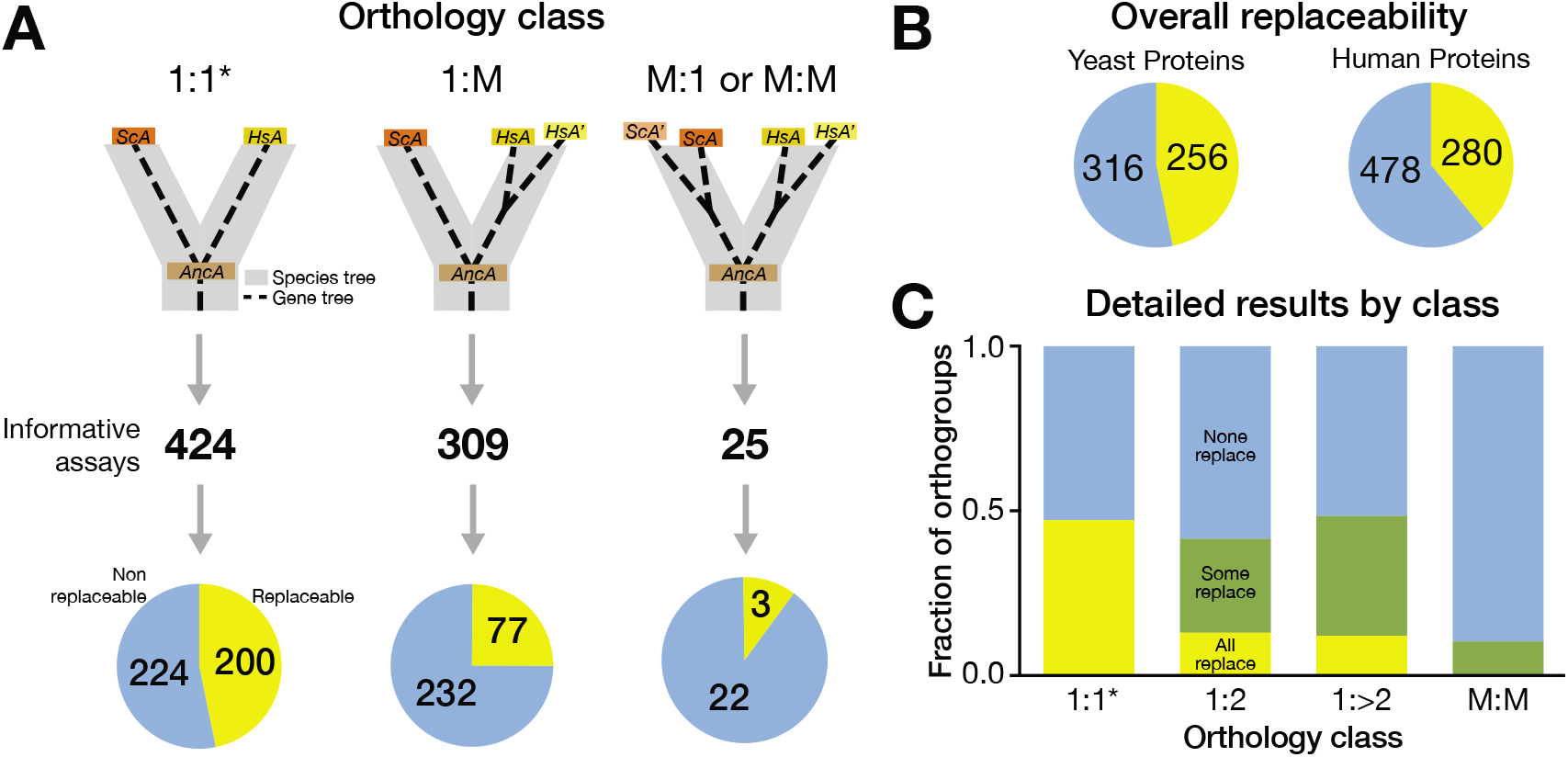
Distribution of replaceability across orthology classes. Only rarely did all human co-orthologs in one orthogroup replace. Rather, a family of human proteins typically had one or a few replaceable members or none at all. **(A)** Previously, systematic replacement of essential yeast genes with 1:1 orthologs (1 yeast to 1 human) demonstrated nearly 50% replaceability of essential yeast genes [18]. Here, we tested the replaceability of essential yeast genes with their human counterparts that have acquired lineage specific duplications in either yeast or human lineage. Of the 309 informative assays obtained in 1:M class (1 yeast to 2 or more human co-orthologs), 77 human genes replaced their yeast equivalents where as 232 did not. Of the 25 informative assays obtained in M:1 and M:M class (>=2 yeast to 1 or more human co-orthologs), 3 human genes replaced their yeast equivalents where as 22 did not. **(B)** Combining our previous replaceability assays [18] with the assays done in this study, we have identified 254 essential yeast genes that are functionally replaceable by their human counterparts and 302 that are not. From the perspective of human proteins, 280 replace their yeast versions while 478 do not. Summary of all the humanized yeast assays performed this far. **(C)** Nearly half of the essential yeast genes belonging to 1:1 orthology class were replaceable by their human equivalents (yellow). The distribution of essential yeast genes replaced by at least one human ortholog in the 1:M (both 1:2 and 1:>2 combined) closely matched the 1:1 results. These yeast genes were rarely replaced by all human co-orthologs in an orthogroup (yellow), with the majority of replaceability falling in the differentially replaceable set (green). M:M yeast orthologs were rarely replaceable by any human ortholog, with only three (of 25 tested) human genes replacing three separate yeast orthologs. * indicates data from [18].

Of the 131 yeast genes for which at least one human ortholog yielded an informative assay, 53 (~40%) had at least one human ortholog that replaced, while 78 were not replaceable by any tested human ortholog (**Figures 2C, S3, and S4, Supplementary File 1**). Of the orthogroups with at least one complementing human gene, the majority (34/53 groups, or 64%) showed differential replaceability while all human co-orthologs in a group replaced only in a few cases (12/53 or ~23%) (**Figure 2C, S3 and S4, Supplementary File 1**). In marked contrast, the great majority of human genes belonging to the M:M class were not able to replace their yeast orthologs, with only three human genes complementing in these cases (**Figures 2C and S4C, Supplementary File 1**).

### Computational analysis of trends governing replaceability

Because the ability of human co-orthologs to functionally replace their singleton yeast orthologs was generally differential within expanded gene families, we sought to identify characteristic features governing selective replaceability of specific human genes in expanded orthogroups. To that end, we assembled and/or calculated a number of quantitative properties for all genes and ortholog pairs. These properties include direct properties of the genes in an orthogroup (e.g. protein length, codon usage bias), as well as comparative properties between co-orthologs (e.g. sequence similarity, length difference). We could then assess each property for its ability to explain replaceability as the area under a receiver-operator characteristic (ROC) curve, treating each feature as an individual classifier, in a similar manner to the previous 1:1 ortholog study [18]. To simplify pairwise comparisons, we considered median values of co-orthologs within each orthogroup that had the same replaceability status *(i.e.* were both replaceable or not) (**Figure S5**). Of the 378 testable 1:M pairs, the majority (104 orthogroups) belonged to a group that had only two human co-orthologs (referred to as the 1:2 class) (**Figure S1**). The remaining orthogroups with more than two human co-ortholog members were dubbed the 1:>2 class. Owing to this disparity, we chose to consider these two groups separately in subsequent analyses.

### Differential replaceability in 1:2 orthogroups is predicted by co-ortholog divergence and mRNA expression specificity

In the 104 1:2 ortholog groups, we obtained informative results for 172/208 human genes (95/104 yeast genes) (**Figures S1 and S3, Supplementary File 1**). 45 (26%) of the tested human genes functionally replaced the yeast gene, while 127 (74%) could not (36 human genes were either not tested or not informative). 77 groups were completely tested *(i.e.* both human co-orthologs were assayed and yielded informative results), 18 groups had only a single tested human gene, and nine had no informative assay results. From the yeast genes’ perspectives, 35/95 (37%) of those assayed were replaceable by at least one human ortholog.

Computational analysis of the 1:2 orthogroups, revealed several explanatory features (**Figure 3A**). Most striking was the relative divergence of the human co-orthologs from each other and from the yeast ortholog. In particular, the highest predictive power (measured as area under the ROC curve, AUC) was seen for the InParanoid ortholog rank (Hs_OrthoRank), which ranks the human co-orthologs from *more-diverged* (MDO) to *least-diverged* (LDO) from the yeast counterpart, and ortholog score (Hs_OrthoScore, a score nearer to 1 being nearer to the yeast ortholog) (**Figure 3A**). Of the 77 completely tested 1:2 orthogroups, there were 22 groups where the two human genes displayed differential ability to replace *(i.e.* one human gene replaced while the other did not). These groups are particularly interesting because they provide an opportunity to investigate properties that distinguish replaceable and non-replaceable co-orthologs within an orthogroup. To assess this difference, we performed ROC analysis for each feature specifically on the set of differentially replaceable 1:2 orthogroups (**Figure 3B**). Notably, the relative co-ortholog divergence remained among the strongest predictors in these orthogroups, with the least-diverged of the two human co-orthologs considerably more likely to replace (**Figure 3B**). Indeed, in this differentially replaceable set, 86% of the replaceable human genes are also the LDO in their respective orthogroups (a trend similar to that observed by Hamza and colleagues [19]). Further, when we compared the divergence of the differentially replaceable set (‘one replaces’) with the ‘both replace’ and ‘none replace’ sets, the average InParanoid ortholog scores of the MDO were similarly low (0.50 and 0.51, respectively), whereas the MDO in the ‘both replaceable’ class is significantly more similar to the LDO (0.66, p-value <0.05) (**Figure 4A**). The mean sequence identities between human and yeast orthologs for the ‘both replace’ and ‘one replaces’ classes are not significantly different, while the ‘none replace’ class has a modestly, though significantly lower mean identity to the yeast ortholog (**Figure 4B**). Thus, while human co-orthologs in the three complementation classes have diverged from their yeast ortholog to a similar extent, those in the ‘both replace’ class have maintained similarity to each other and are thus more likely to both replace (**Figure 4C-E**).

**Figure 3.**
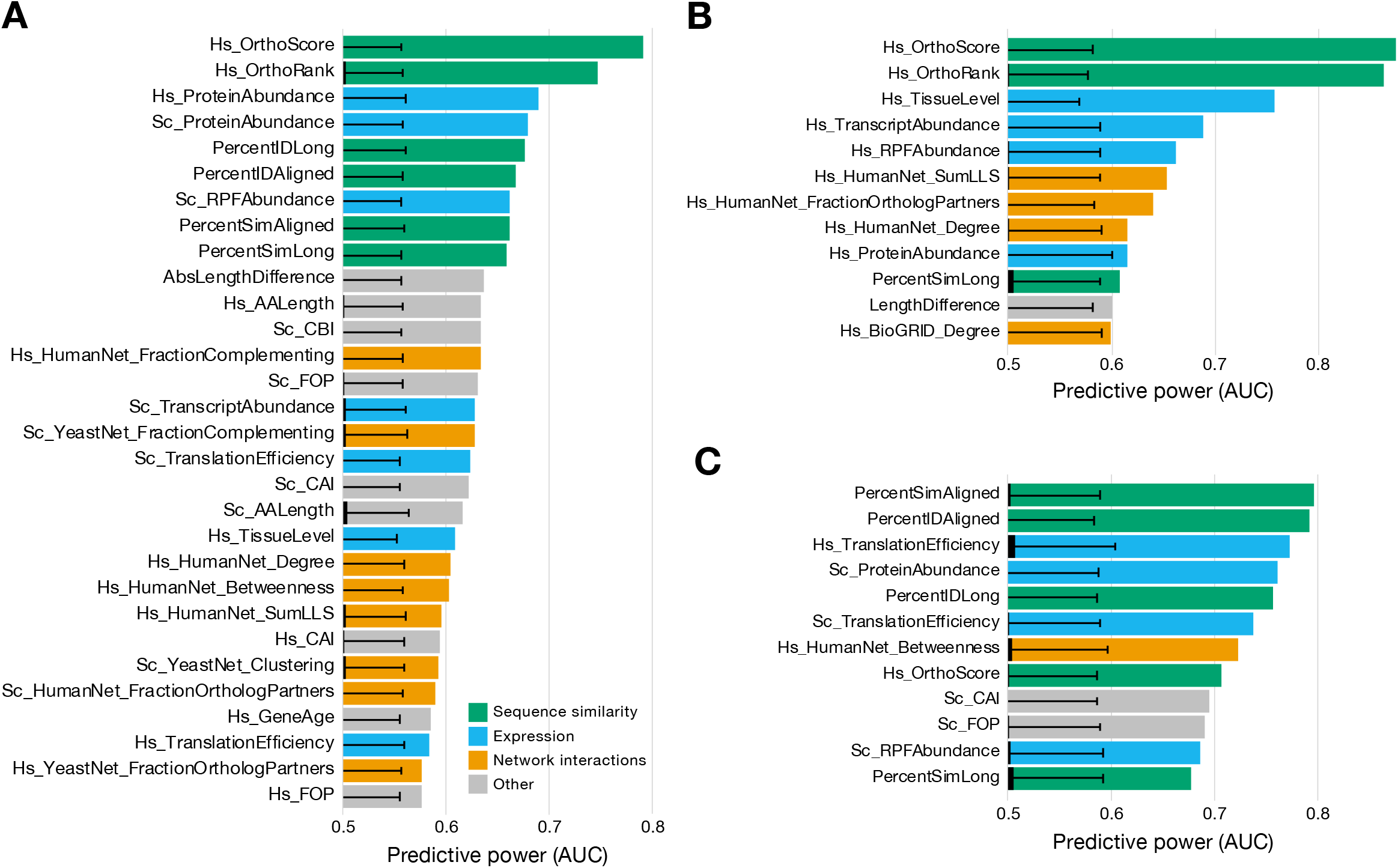
Replaceability of 1:2 orthologs is explained largely by relative divergence of human co-orthologs. **(A)** Area under the ROC curve (AUC) for the top 30 predictive features for median-collapsed 1:2 orthogroups are shown. The top two predictive features (HsOrthoScore and HsOrthoRank) for this class indicate that replaceability is driven largely by the nearness of the replaceable human co-ortholog to the yeast gene relative to a non-replaceable co-ortholog (*i.e.* most 1:2 orthogroups have only one replaceable human co-ortholog, and it is almost always the least-diverged one). **(B)** The observed trend is even more strongly demonstrated when analysis is restricted to the specific set of 1:2 orthogroups that display differential replaceability (*i.e.* one human co-ortholog replaces and the other does not), with AUCs nearing 0.9. The extent of tissue-specific expression also becomes significantly predictive in this set, indicating that human co-orthologs that are more broadly expressed are more likely to replace than their more tissue-specifically expressed co-ortholog. **(C)** More-diverged human co-orthologs in a 1:2 pair do replace in several cases (mostly those where both co-orthologs replace). When restricting analysis to this set, it is apparent that the most predictive property is sequence similarity to the yeast ortholog, along with proteins that are translated efficiently and are typically in high abundance. Black overlapping bars indicate mean and error bars indicate standard deviation for 1000 shuffled AUC calculations for each feature.

An additional significantly predictive feature appearing in the differentially replaceable 1:2 class pertains to the tissue-specific expression of the human orthologs. Specifically, co-orthologs with more widespread (less tissue-specific) expression were more likely to replace (**Figure 3B**). This feature is particularly predictive of the differentially replaceable set, but was not a strong feature for the full 1:2 orthogroup results (**Figure 3A**).

### More diverged yet replaceable 1:2 orthologs are highly expressed and more similar to their yeast counterparts

Despite the fact that in the majority of the 1:2 cases the least diverged human co-ortholog replaced the corresponding yeast gene, there were several cases wherein the more-diverged co-ortholog was replaceable. Of 81 informative MDO assays, 13 human MDOs replaced. To determine what drives these replacements, we applied our ROC analysis scheme to the set, not restricting it to completely assayed groups. For these genes, the ability to complement is dominated by sequence similarity to the yeast ortholog as well as protein abundance features (**Figure 4C**). These results are in line with previous observations that human co-orthologs that both replace are more similar than those where only one or none replaces, but also suggests that these highly diverged co-orthologs retain high expression levels. Thus, some highly diverged human co-orthologs still seem to maintain ancestral functionality, irrespective of the other co-ortholog’s function.

**Figure 4.**
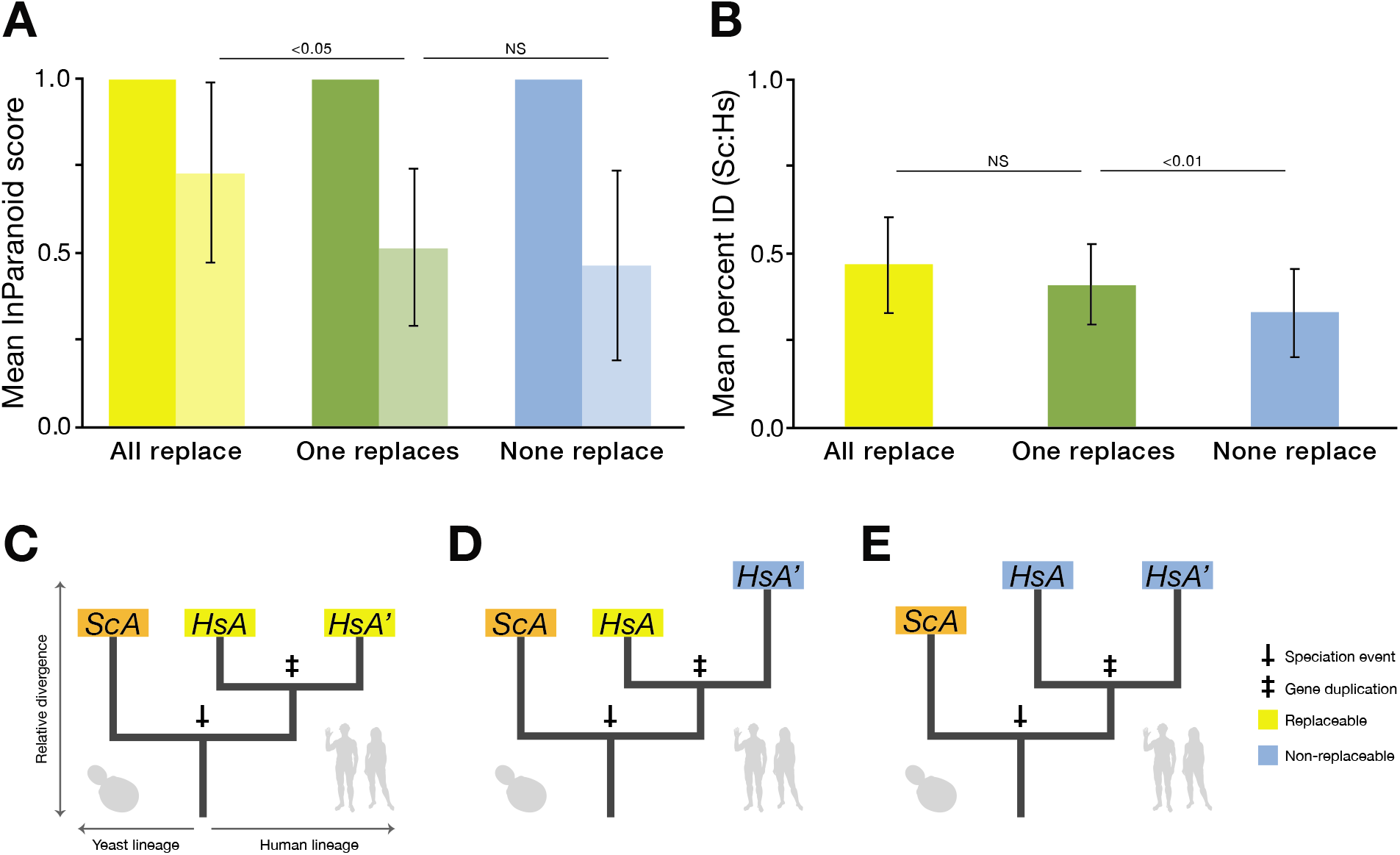
Replaceability is explained by relative divergence of 1:2 human co-orthologs from each other and their yeast ortholog. **(A)** The average ortholog score of the more-diverged (lighter color) 1:2 co-ortholog is more similar to the less-diverged co-ortholog for co-ortholog pairs that both replace than pairs where only one or neither of the co-orthologs replace. **(B)** The average percent amino acid identity for 1:2 co-orthologs are not significantly different between the ‘all replace’ (yellow) and ‘one replaces’ (green) class, while the ‘none replace’ class is slightly, but significantly, lower than either. Error bars indicate standard deviation. **(C-E)** Phylogenetic models depicting gene trees for generic 1:2 orthogroups in the various replaceability classes. In **(C)**, the human co-orthologs belong to the ‘both replace’ class and are thus less diverged from each other but are on average similarly diverged from the yeast ortholog as the ‘one replaces’ class **(D)**, where the co-ortholog more closely related to the yeast gene is the replaceable one. In **(E)**, the ‘none-replace’ co-orthologs are on average more diverged from the yeast gene.

### Human co-ortholog replaceability in 1:>2 orthogroups is marked by conserved interactions and subcellular localization

In the case of 1:>2 orthogroups, 137/170 human genes were successfully assayed, 32 of which replaced the yeast ortholog while 105 did not (23% replaceable). There were 23 1:>2 groups for which we obtained informative results for all human co-orthologs, represented by 78 pairwise tests. From the yeast perspective, 18/36 (50%) of yeast genes with at least one informative test were replaceable by at least one human co-ortholog. Six of those were replaceable by only one co-ortholog, and only two were replaceable by all co-orthologs (**Supplementary File 1**).

As orthogroups with more than two human members had differing rates of replaceability within them and not all human genes could be assayed, we sought to simplify the analysis of features that differentiate between co-orthologs that can replace the yeast ortholog and those that cannot (**Figure 5A**). We thus considered median properties of replaceable vs. non-replaceable genes in each 1:M orthogroup (**Figure S5**). Unlike for 1:2 orthogroups, (**Figure 5A**) while sequence similarity appears near the most explanatory features, the ability of 1:>2 orthologs to replace was most strongly marked by cellular context. The dominant predictive features included the fraction of conserved protein-protein interactions along with their centrality in their respective functional interaction networks and the number of interactions in those networks, albeit to a lesser extent. Specifically, human co-orthologs that have maintained a higher fraction of orthologous interaction partners relative to the yeast ortholog were more likely to replace. Further, human co-orthologs with relatively higher centrality in a functional network were more likely to be replaceable. Thus, in the case of highly expanded gene families, paralogs that maintain ancestral protein contacts and centrality in their interaction networks tend to be more replaceable (**Figure 5A**), consistent with previous observations that centrally placed proteins in interaction networks tend to carry out crucial cellular functions [31] that have been retained over vast evolutionary timescales.

**Figure 5.**
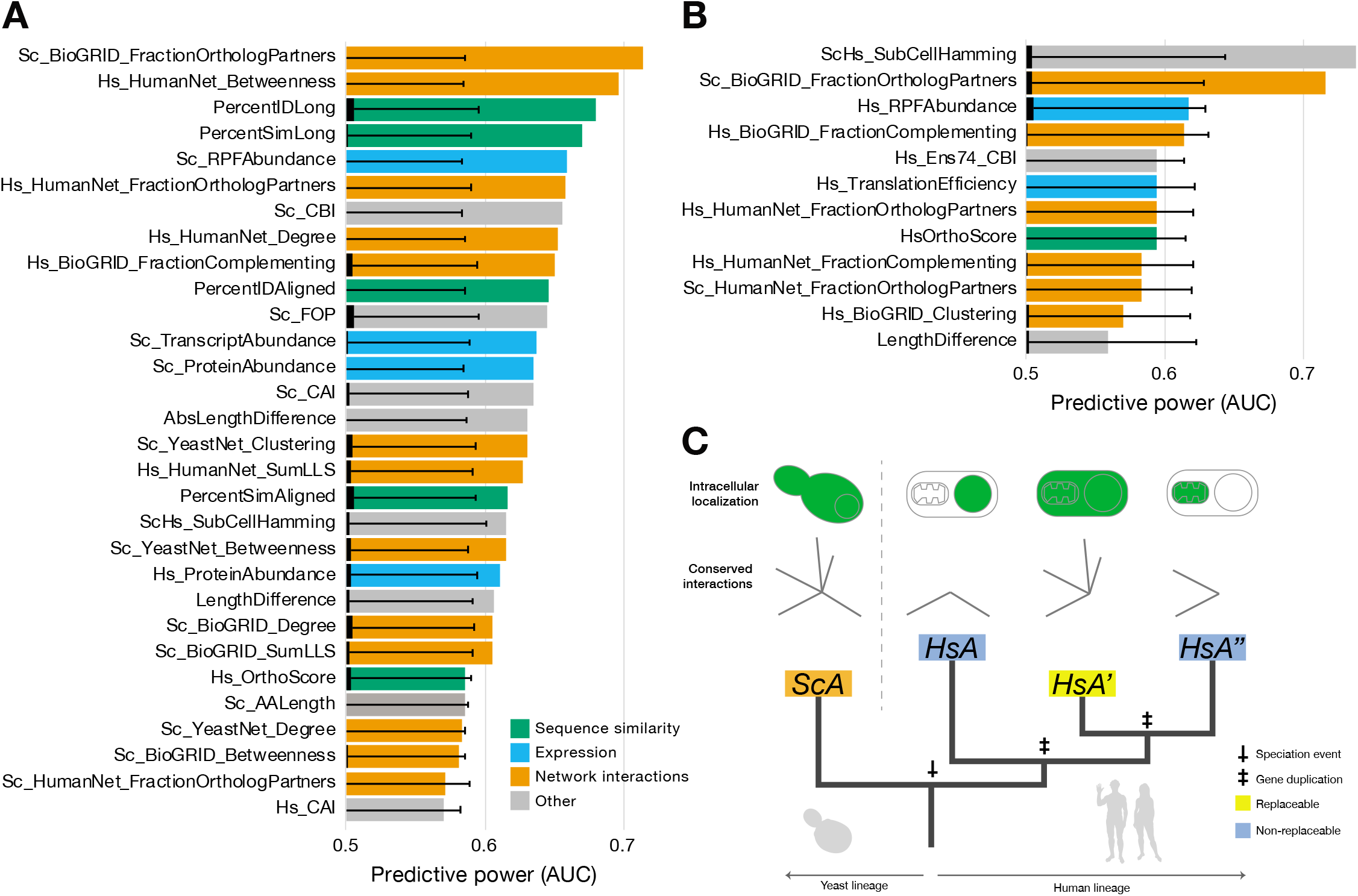
Replaceable 1:>2 human co-orthologs retain orthologous interaction partners and are more central in interaction networks. **(A)** AUCs of the top 30 predictive features for median-collapsed informative 1:>2 co-ortholog pairs. The top two most significant features demonstrate the importance of network context in retaining ancestral functions. Specifically, human co-orthologs in highly expanded gene families that have retained a higher fraction of orthologous protein interactions partners with their yeast ortholog (FractionOrthologPartners) are more likely to replace, as well as those that maintain higher centrality in functional interaction networks (Betweenness). **(B)** Similar to the 1:2 case, we further restricted our analysis to a subset of median-collapsed 1:>2 orthogroups that had both replaceable and non-replaceable human co-orthologs. In this set, the subcellular localization of the human proteins appears to be predictive, in that more broadly localized co-orthologs are more likely to replace than their more organellar-specific co-orthologs. While this AUC is not significant (indicated by being just less than two standard deviations off the mean), it is an obvious trend and does overtake fraction of orthologous partners as the highest performing AUC for this set. **(C)** Phylogenetic model of a generic 1:M orthogroup, showing that the replaceable human co-ortholog has retained more orthologous interactions in a network, and is localized in a similar manner to the yeast ortholog. For AUC bar plots, black overlapping bars indicate mean and error bars indicate standard deviation for 1000 shuffled AUC calculations for each feature.

We further sought to identify features that distinguished replaceable co-orthologs from non-replaceable ones within the same orthogroup for the 1:>2 set. We again considered median features of both replaceable versus non-replaceable co-orthologs within those 1:>2 orthogroups that showed differential replaceability. No features showed an AUC more than two standard deviations above the mean of permutation tests, likely due to the small size of this specific orthogroup set. Nonetheless, the strongest trend observed was for replaceable co-orthologs to be localized in more similar subcellular compartments [32] in human cells as the yeast ortholog(s) (**Figure 5B**). Our observations are consistent with a model that at least one co-ortholog in an expanded human gene family will tend to retain essential ancestral functionality by maintaining ancestral cellular localization and ancestral centrality in the functional interaction network of the cell (**Figure 5C**).

### Simulations suggest more diverged duplicates are less likely to bind their ancestral interaction partners

To assess the generality of our observations, we performed *in silico* simulations of functional divergence in a duplicated gene family. We analyzed a small heterodimeric protein complex consisting of A and B subunits (the SMT3-UBC9 protein complex [18,33]), where the subunit B was duplicated *in silico* to yield A, B, and B’. Using binding of A to B and/or B’ as a proxy for functionality, we carried out evolutionary simulations of molecular structural divergence (considering all atom models using the Rosetta molecular modeling suite [34,35]. As described in [36], we examined functional replaceability across five different selection scenarios. All of the simulations assumed that selection acts on the stability of each subunit but differed in how they imposed selective pressure on binding. We ran 100 replicates for each of the five selection schemes, and quantified replaceability of each of the subunits (measured as continued ability to bind to the ancestral partner) over sequence divergence (**Figure 6A, B**). Notably, selection for A to bind both B and B’ results in a continued ability for either B or B’ to bind the ancestral variant A, whereas application of diversifying selection to prevent binding to B’ results in a rapid decay in the ability of B’ to bind A (**Figure 6B**).

**Figure 6.**
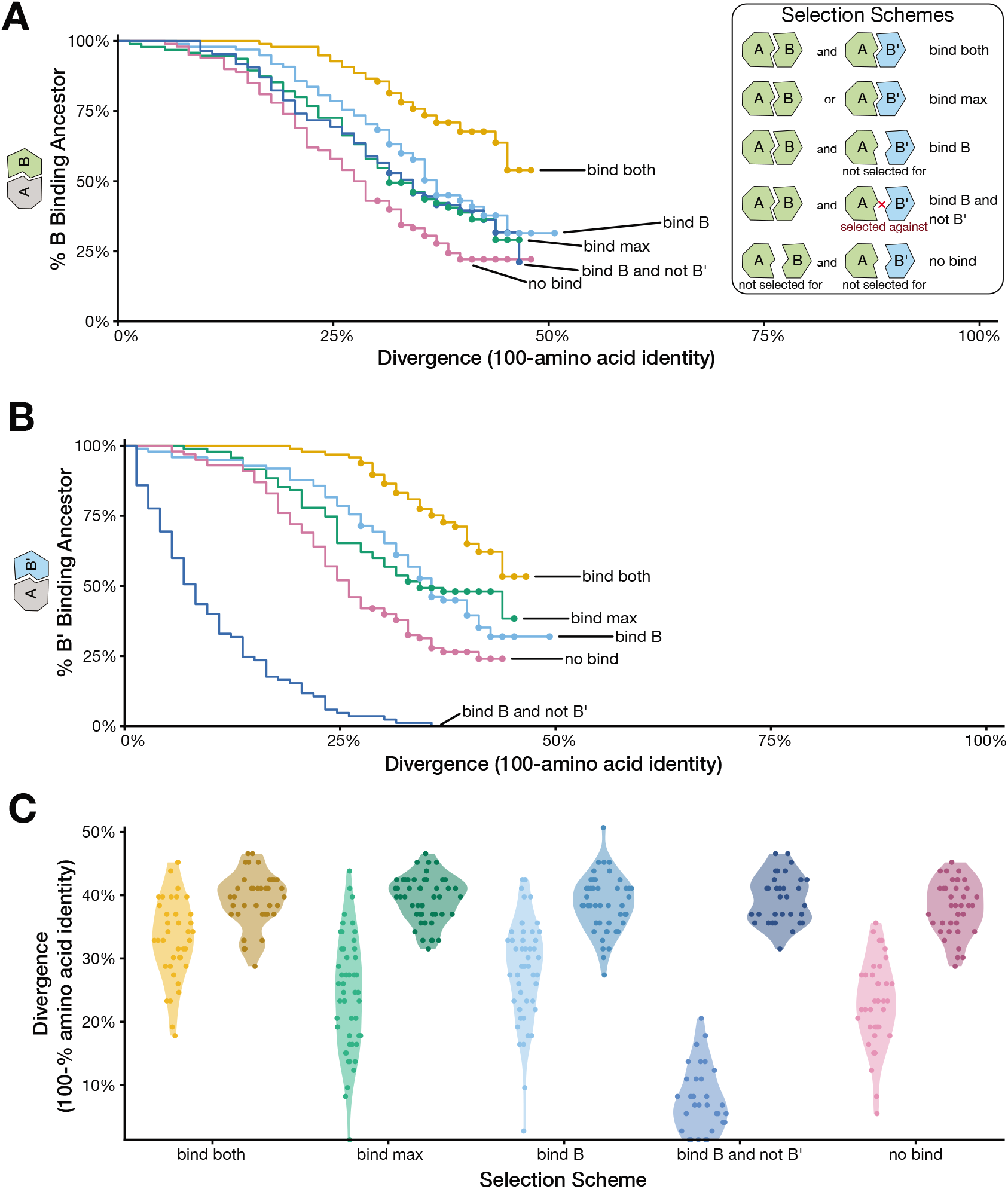
Simulated protein evolution suggests diverged duplicates are less likely to bind their ancestral interaction partner. Five different types of selection scenarios were considered for a heterodimeric protein complex, considering the effects of amino acid substitutions using the Rosetta molecular modeling platform. (1) AB and AB’ (Bind both). (2) AB or AB’, and only the most stable binding interface is considered (Bind max). (3) AB but AB’ was not enforced (Bind B). (4) AB and AB’ is selected against (Bind B and not B’). (5) Neither AB or AB’ is selected for (No bind). Figure adapted from [36]. **(A)** Percent of simulations where an evolved B subunit has the ability to bind the ancestor to A. Divergence is measured by the amount B has diverged from the ancestor of B. **(B)** Percent of simulations where an evolved B’ subunit has the ability to bind the ancestor to A. Divergence is measured by the amount B’ has diverged from the ancestor of B. **(C)** Percent divergence of duplicates when only B or B’ is able to bind the ancestor A. Lighter hues denote duplicate that is able to bind the ancestor of A, and darker hues denote the non-binding duplicate.

We then looked across our simulated lineages at those cases where one of the duplicates functionally replaces *(i.e.* binds to the ancestor) and the other does not. We found a systematic pattern, across all five selection schemes, that the non-binding duplicate tends to be the more diverged one (**Figure 6C**; all pairwise comparisons within selection schemes are significant, *p* < 0.001 paired t-test). These results mirror our experimental humanization findings for the relative divergence of 1:2 human co-orthologs that the replaceable duplicate tends to be less diverged than the non-replaceable duplicate (**Figure 4**).

## Conclusions

By extending the scope of our systematic yeast humanization assays to include those yeast genes that have more than one human ortholog, we successfully added 309 human genes to our tested set (a 73% increase). We have therefore greatly expanded the set of human genes that can now be functionally studied in the simplified unicellular eukaryotic context of budding yeast by adding 80 human genes (56 novel to this dataset) to those that can successfully replace their yeast ortholog. Overall, we found that yeast genes with duplicated human orthologs can be replaced by at least one co-ortholog at a slightly lower rate (40%) than 1:1 orthologs [18]. Of those that could be replaced, the clear majority were replaceable only by one or two co-orthologs, rather than being broadly replaceable by many human genes in the same family. This pattern of replaceability between and within orthogroups supports the long-held hypothesis that orthologs between species will retain ancestral functions at a higher rate than paralogs within a species [10,11].

After testing many properties of the gene families for their ability to explain replaceability, our analysis revealed divergent patterns across groups that have two (1:2) or more than two human (1:>2) human co-ortholog members. In the case of the 1:2 class, the top predictors were dominated by features that capture divergence from the yeast ortholog, in particular we observed that the less-diverged variant tended to strongly be the replaceable one. This observation was supported by computational simulations of protein divergence after duplication, where the least diverged of the duplicates retained ancestral binding ability in most cases. In addition to sequence divergence, we also observed a significant trend of replaceable 1:2 human co-orthologs to have broader, less tissue-specific expression. Somewhat rarely, the more-diverged human co-ortholog could replace, and these tended to be more similar to their yeast counterpart and expressed at a higher level perhaps due to retaining important ancestral functions. In the case of the 1:>2 set, replaceability was marked largely by network-based properties of the genes. Replaceable co-orthologs in this set seem to have retained more ancestral interaction partners as well as higher centrality, even across a billion years of divergent evolution, likely indicative of their functional importance. While not significant by our criteria, we also observed replaceable 1:>2 co-orthologs being expressed in similar subcellular compartments to their corresponding yeast gene, highlighting a trend to maintain their ancestral localization.

Overall, our extended set of humanization assays and their analysis reveal principles of functional divergence in paralogs and orthologs. We observed a strong tendency for orthogroups to exhibit only one or a few swappable human genes rather than many. We also extended our observation that network centrality and interaction properties aid in determining how ancestral gene function across orthologs is retained over deep evolutionary timescales.

Such assays help to advance our understanding of duplicate gene evolution in the billion years since yeast and humans diverged [37] and add powerful reagents to study myriad human processes and develop therapies in a simpler eukaryotic surrogate.

## Materials and methods

### Identifying orthologs

Orthologs were calculated with a local installation of InParanoid [25], using UniProt proteomes for the two species (downloaded November 2014). The InParanoid script was modified to use a more recent version (2.2.25+) of the BLAST+ suite of programs [38].

InParanoid identifies orthogroups between two species by first performing an all-vs-all BLAST search between the two species to identify bidirectional best-hits (BBH). Each proteome is then subjected to an all-vs-all BLAST against itself to identify within-species homologs. The BBH pairs are used as seed matches and any within-species genes from the self-BLAST that are at least as close to the gene of interest as its BBH in the other species is added to the ortholog group and termed an ‘in-paralog’ or co-ortholog. We used the ‘table’ output of InParanoid to identify orthogroup classes as follows: Those groups with one listed yeast gene and two human genes were dubbed one to two (1:2) orthologs, while those with one yeast gene and more than two human genes were identified as one to more-than-two (1:>2) orthologs. Together, these two sets make up the one to many (1:M) ortholog class.

### ORFeome cloning

Human genes were obtained from the human ORFeome collection [26]. The ORFeome comprises a collection of *E. coli* strains, each harboring a plasmid encoding a single human gene in a Gateway ‘entry’ vector. Sequences cloned in the entry vectors are flanked by attL sites. To create expression vectors, each human entry vector was isolated from *E. coli*, added to a Gateway LR reaction with a Gateway ‘destination’ vector and transformed into competent *E. coli* to obtain expression clones [39]. As ORFeome clones lack stop codons, we modified the Advanced Yeast Gateway kit [28] pAG416-GPD-ccdB destination vector which does not encode a stop codon immediately outside of the cloning region resulting in a ~60 amino acid tail being added to any protein expressed from it. We thus mutagenized the vector downstream of the cloning region to introduce a stop codon, shortening the tail to six amino acids (this plasmid is termed pAG416-GPD-ccdB+6Stop) [18]. Entry and expression clones were verified by sequencing into the gene sequence from the upstream and downstream region of the plasmid.

### MGC cloning

For human genes not available in the ORFeome, we obtained clones from the Mammalian Gene Collection [27], a collection of sequence-verified human cDNA sequences. To obtain entry vectors for these genes, we designed primers for each gene that would amplify the coding sequence while adding attB sites to either end of the human gene and performed PCR using the MGC plasmid as template. PCR products were combined with plasmid pDONR221 in a Gateway BP reaction [39] and transformed to *E. coli* to obtain entry clones. The entry clones were then combined in a Gateway LR reaction with p416-GPD-ccdB+6Stop and transformed to *E. coli* to obtain expression clones. Each entry and expression clone was verified by sequencing the clone boundaries at each end of the gene.

### Functional replaceability assays

Yeast strains were grown in a 96-well format in YPD medium supplemented with G418 [200 *μ*g/ml]. The strains were transformed with matched expression clones or empty control vectors and selected on minimal medium lacking uracil. Complementation assays were performed as follows:

#### Temperature-sensitive assays

Each of the strains in the temperature sensitive (TS) strain collection [30] encode a yeast protein with a mutation that allows growth at permissive temperatures of 22-26°C, but not at the restrictive temperature of 35-37°C. We therefore identified human genes capable of rescuing growth of the mutant at restrictive temperature on selective plates. Each strain was transformed and assayed separately with either the human gene-expressing vector or the empty vector control by growing transformed strains in the following manner:

-Ura dextrose medium at the permissive temperature (26°C), serving as a control for transformation efficiency and/or toxicity since both the yeast and the human gene are expressed.
-Ura dextrose medium at the restrictive temperature (37°C), testing for human gene functional replacement under conditions in which the corresponding yeast gene is non-functional.

#### Magic Marker assays

Strains in the Magic Marker collection (obtained from ATCC) are heterozygous diploids, each harboring one allele of a yeast gene knocked out by replacement with the KanMX kanamycin-resistance cassette, allowing for selection on G418 [29]. We transformed human expression clones or an empty control vector into appropriate strains and selected on-Ura G418 medium in a 96-well format. Transformants were then re-plated on GNA-rich pre-sporulation medium containing G418 and 50 mg/L histidine. Individual colonies were then inoculated in liquid sporulation medium containing 0.1% potassium acetate, 0.005% Zinc acetate, and incubated with vigorous shaking at 26°C for 3-5 days, after which sporulation efficiency was estimated by microscopy, the mixture then re-suspended in water and equally plated on two assay conditions:

1. -G418 Magic Marker dextrose medium (−His −Arg −Leu +Can −Ura), incubated at 30°C. The haploid spores that carry the wild-type allele grow in this medium providing us with the control for sporulation efficiency. This condition also assays for toxicity of the human gene if the haploid spores fail to grow.
2. +G418 Magic Marker dextrose medium (−His −Arg −Leu +Can −Ura) containing 200*μ*g/ml G418. In the absence of the human gene (as for control transformants), the resulting haploid knockout strain is expected not to grow, providing an assay for replaceability in strains expressing the human gene. Cases with approximately equal numbers of colonies growing in the absence or presence of G418 were considered functional replacements. For cases with ambiguous growth (marked by moderate numbers of isolated colonies growing on the +G418 medium relative to −G418 medium), we screened varying quantities of the sporulation mixtures.

Positive assays were verified independently. Individual colonies were isolated from selective plates and were assayed for growth defects on YPD or Magic Marker medium+ G418 (**Figures 1B, S3 and S4**). After growth on YPD+G418, each strain was spotted on 5FOA agar to test plasmid dependency.

### Tetrad dissection and plasmid loss assays

For human gene replaceability assays performed in the yeast heterozygous knockout Magic Marker collection that were ambiguous in our large-scale screen, we performed tetrad dissections to more clearly test for complementation (**Figure S2**). In total, 33 human genes were assayed and analyzed. We transformed each human expression clone or empty vector control into the appropriate yeast strains and selected on SC–Ura + G418 (200*μ*g/ml) to select for the human gene expression vector (CEN, Ura+) and yeast gene knockout (KanMX marker) simultaneously. Transformants were then plated on GNA-rich pre-sporulation media containing G418 (200*μ*g/ml). Individually isolated colonies were inoculated into liquid sporulation medium containing 0.1% potassium acetate, 0.005% Zinc acetate and were incubated with vigorous shaking at 25°C for 3-5 days. Following this, sporulation efficiency was estimated by microscopy, and successful sporulations were subjected to tetrad analysis. 15-20 μL of each sporulation was digested with an equal volume of Zymolyase (5 mg/ml stock) for 30-45 min to remove the ascus coats. The digestions were diluted 1:1 with sterile water after which 20-30 μL of the zymolyase-treated spores were carefully applied to a tilted YPD plate using a pipette, allowing the droplet of cell suspension to gently run down the agar surface. The plates were dried and visualized on the tetrad dissection microscope. For each human gene, a minimum of 5 tetrads were dissected. Each tetrad dissected plate was incubated at 30°C for 2-3 days. Plates containing 3 or more sets of 4 viable spores were selected and replica plated both on 5-FOA (for plasmid counter-selection) and YPD+G418 (for yeast null allele selection). A successful complementation consists of 2:2 segregation with survival on YPD+G418 and failure to grow on 5-FOA (**Figure S2**). Tetrads failing to undergo 2:2 segregation were classified as non-complementing (summarized in **Supplementary File 1**). We subsequently performed quantitative growth assays (in triplicate) on tetrads passing the 5FOA/G418 segregation test. Each humanized tetrad was grown in 3 different media conditions, YPD, YPD+G418 and SC-URA. Each medium-specific growth profile (shown in one of the conditions as in **Figures S3 and S4**) was analyzed and quantified to detect any growth defects in yeast. 14 out of 33 human genes assayed in this manner showed functional replaceability of the yeast gene function (Supplementary File 1).

### Isolating haploid humanized yeast strains and quantitative growth curve assays

Select yeast heterozygous knockout magic marker strains carrying human gene expression vectors (CEN, Ura+) showing functional replaceability were sporulated and plated on independent petri plates to obtain single colonies. Each colony was tested for plasmid loss in the presence of 5FOA. Colonies that resisted plasmid loss were isolated and subjected to further analysis to quantitatively measure their growth rates. Yeast strains were either pre-cultured in liquid YPD + G418 (200*μ*g/ml) or −Ura Dextrose selective medium + G418 (200*μ*g/ml) for 2 hrs or overnight respectively. Each culture was diluted in YPD or −Ura Dextrose medium to an OD of ~0.1 in 100 or 150 μL total volume in a 96-well plate. Plates were incubated in a Synergy H1 shaking incubating spectrophotometer (BioTek), measuring the optical density every 15 min over 48 hr. Growth curves were performed in triplicate for each strain by splitting the pre-culture into three independent cultures for each 48–60 hr time course (**Figures S3A, S4A and S4C**).

In the case of temperature-sensitive humanized yeast strains, the growth assays were performed first at permissive temperatures 25–26°C for 48-60 hr time course. These growth assays were largely identical to the empty vector transformed yeast strains. The strains where then shifted to a restrictive temperature of 37°C and similarly repeated the growth assay for a 48–60 hr time course (**Figures S3B and S4B**). Growth curves were performed in triplicate for each strain by inoculating cells from the same pre-culture into three independent cultures.

For computational analyses of the trends underlying replaceability, we computed a diverse set of features (**Supplementary File 3**), based in part on those from Kachroo *et al.* [18], as follows:

### Sequence properties

Sequence features for human genes obtained from the human ORFeome [26] were calculated using the ORFeome provided fasta file for the sequences, downloaded from http://horfdb.dfci.harvard.edu/hv7/docs/human_orfeome71.tar.gz. For clones not obtained from the human ORFeome collection, we calculated sequence features using the longest annotated transcript or its translation from Ensembl version 74, available at http://Dec2013.archive.ensembl.org/index.html. To analyze protein features, we translated the above nucleotide sequence for each gene using the standard genetic code. The following protein sequence features were considered:

#### Sequence length

Sc_Length

Hs_Length

Sc-Hs_LengthDifference

Sc-Hs_AbsLengthDifference

Length was calculated as the count of amino acids in the protein. LengthDifference was calculated as the length of the human ortholog protein subtracted from the length of the yeast ortholog. AbsLengthDifference is the absolute value of LengthDifference.

#### Sequence similarity

Sc-Hs_PercentIDLongest

Sc-Hs_PercentIDAligned

Sc-Hs_PercentSimLongest

Sc-Hs_PercentSimAligned

Hs_Orthoscore

Hs_OrthoRank

Orthologous pairs were first identified by InParanoid [25]. Global alignments for all pre-identified ortholog pairs were then calculated using NWalign (http://zhanglab.ccmb.med.umich.edu/NW-align/) with BLOSUM62 and gap open penalty of −11 and extension −1. Identity and similarity were calculated as the fraction of identical amino acids and amino acids with positive BLOSUM score in the alignment, respectively. ‘Longest’ refers to amino acid identity or similarity calculated as a fraction of the longer of the two orthologs. ‘Aligned’ refers to calculating identity or similarity calculated as a fraction of the length of the aligned region of the sequences. OrthoScore and OrthoRank refer to the scores assigned by InParanoid. For each orthogroup, InParanoid calculates a confidence score between 0 and 1 for each in-paralog that represents how similar it is to the seed ortholog. Rank is simply a ranked ordering of the inparalogs of a group based on their orthoscore, with 1 being the least diverged ortholog and higher values being further away from the seed.

#### Codon usage

Sc_CAI

Sc_CBI

Sc_FOP

Hs_CAI

Hs_CBI

Hs_FOP

Calculated from the amino acid sequences using CodonW (http://sourceforge.net/projects/codonw/). Properties for the human genes were calculated using the yeast codon optimality table, as a measure of closeness or divergence from yeast optimality.

### Network properties

Network features were calculated using custom Python scripts, typically utilizing the package ‘networkx’ (available from http://networkx.github.io/documentation/latest/download.html). When applicable (e.g. HumanNet, YeastNet), provided weights of interactions were taken into account when calculating these features. Otherwise, a default weight of 1.0 was used. Network features were defined as follows: Degree represents the count of interaction partners for a node in a given network. Betweenness represents network centrality, a measure of how central in a network a given node is, calculated as the number of shortest paths between all node pairs in a network that pass through a given node. Clustering represents the node clustering coefficient, calculated as the fraction of edges that could possibly be present in a node’s neighborhood that are actually present. FractionComplementing is the fraction of interaction partners observed to complement (including results obtained in our 1:1 humanization assays [18]). FractionOrthologPartners is the fraction of a genes interaction partners that have orthologs in the other species, and interact with the gene of interests’ ortholog in the corresponding network (*i.e.* if gene Sc*-A* interacts with Sc-*X*, Sc-*Y*, and Sc-*Z*, and Hs-*A* interacts with Hs*-X* and Hs-*Y*, but not Hs*-Z* (which is a legitimate gene), the FractionOrthologPartners for Sc-A is 2/3 or 0.66). It represents a measure of the degree to which interactions are maintained between orthologs in the two species.

#### BIOGRID

[HslSc]_BIOGRID_[DegreelBetweennesslClusteringlFractionComplementinglFractionOrthologP artners]

Calculated from interactions present in BIOGRID 3.1.93 [40], using only those interactions annotated as ‘physical interactions’.

#### Functional Networks

[HslSc]_*Net_[DegreelBetweennesslClusteringlFractionComplementinglFractionOrthologPartners]

Human and yeast functional gene network features were calculated based on HumanNet [41] and YeastNet [42], respectively. The final sum log-likelihood score reported for each interaction was employed as an edge weight for calculations.

#### Abundance properties

[HslSc]_ProteinAbundance

[HslSc]_T ranscriptAbundance

[HslSc]_RPFAbundance

[HslSc]_TranslationEfficiency

ScHs_SubCellHamming

Protein abundances were used as reported by Kulak *et al.* [43]. TranscriptAbundance, RPF (Ribosome Protected Fragments) Abundance and TranslationEfficiency were calculated from Guo *et al.* (Human) [44] and Ingolia *et al.* (Yeast) [45]. Translation efficiency was calculated as the ratio of RPF reads to mRNA reads. SubCellHamming is a measure of the difference in subcellular localization between a yeast-human ortholog pair, calculated using data from the COMPARTMENTS database ([32], downloaded in February 2017). We utilized the ‘benchmark’ sets to create, for each protein in their respective species, a binary vector of subcellular localization across 11 compartments (Cytoskeleton, Cytosol, Endoplasmic Reticulum, Extracellular space, Golgi apparatus, Lysosome, Mitochondrion, Nucleus, Peroxisome, Plasma membrane) that were common to both human and yeast in the database. Then, for each ortholog pair, we calculated the hamming distance between their respective subcellular localization vectors as the SubCellHamming value.

### Calculating predictive strength of features

The predictive power of each feature was calculated as the area under the receiver-operator characteristic curve (AUC) when treating that feature as an individual classifier. Each feature was sorted in both ascending and descending directions, retaining the direction providing an AUC > 0.5. To assess significance, a permutation procedure was performed as follows: For each feature, the replaceable/non-replaceable status of each ortholog pair was shuffled (retaining the original ratio of replaceable to non-replaceable assignments), and the AUC was calculated. The shuffling procedure was carried out 1,000 times for each feature, and the mean AUC values and their standard deviations reported.

#### Features of expanded orthogroups

In order to avoid overweighting expanded gene families and compensate for uneven sampling, we considered median properties of paralogs in each family as follows:

For each orthogroup, we collapsed all yeast-human ortholog pairs with the same status (replaceable or not) into a single case, with the value of each feature was calculated as the median value of the collapsed ortholog pairs for that feature from the original table. Thus, each 1:M orthogroup is collapsed to be represented by either two pairs (Complement and Non-complement) or one pair (either Complement or Non-complement).

### Simulations

We constructed a simulation of protein evolution where one member of an interacting pair was duplicated. The simulation was initialized with the yeast SMT3-UBC9 complex, a small heterodimeric protein complex, as the resident genotype (PDB: 2EKE) [33]. This complex initially had two subunits, which we refer to as A and B. We duplicated the B subunit and refer to it as B’. Our simulation protocol and setup [36] is based on an accelerated origin—fixation model [18,46]. Here, we further analyzed five of the previously published sets of 100 simulation trajectories [36] (summarized in the **Figure 6A Selection Schemes panel**). Briefly, under the first selection scheme we enforced selection for A to bind B and for A to bind B’ (bind both). In the second scheme selection acts on the ability of A to bind B or A to bind B’ and the maximum stability of those interactions was considered (bind max). In the third scheme the ability of A to bind B was selected for but the ability of A to bind B’ was not (bind B). In the fourth scheme the ability of A to bind B was also selected for but the ability of A to bind B’ was selected against (bind B and not B’). We also implemented a control selection scheme where selection does not act on the ability of A to bind either B or B’ (no bind) We performed 100 replicates of each of these selection schemes.

The percent of simulations where an evolved duplicate has the ability to bind the ancestral partner (**Figure 6A, B**) correspond to Figure S7A, B in [36]. We further analyzed these data by examining instances where only one of the duplicates is able to bind the ancestral partner at the end of the simulation run. We recorded each duplicate’s divergence, defined as the fraction of amino acid positions that were non-identical between the initial and final sequences of a simulation run (**Figure 6C**). For each selection scheme, the distribution of the divergence of duplicates that can bind the ancestral partner was compared to the distribution of divergence of duplicates that cannot bind the ancestral partner with a paired t-test.

## Acknowledgements

The authors would like to thank Charles Boone (University of Toronto) for generously sharing the ts yeast collection and to Maitreya Dunham (University of Washington) for constructive advice. This research was funded by grants to R.K.G. from the American Heart Association Predoctoral fellowship (#18PRE34060258), to A.H.K. from the Natural Sciences and Engineering Research Council (NSERC) of Canada Discovery grant (RGPIN-2018-05089), CRC Tier 2 (NSERC/CRSNG-950-231904) and Canada Foundation for Innovation and Québec Ministère de l’Économie, de la Science et de l’Innovation (#37415), to C.O.W from the National Institutes of Health (R01 GM088344) and National Science Foundation Co-operative Agreement (DBI-0939454 BEACON Center), and to E.M.M. from the Welch Foundation (F-1515) and National Institutes of Health (R35 GM122480).

---

**Figure S1.**
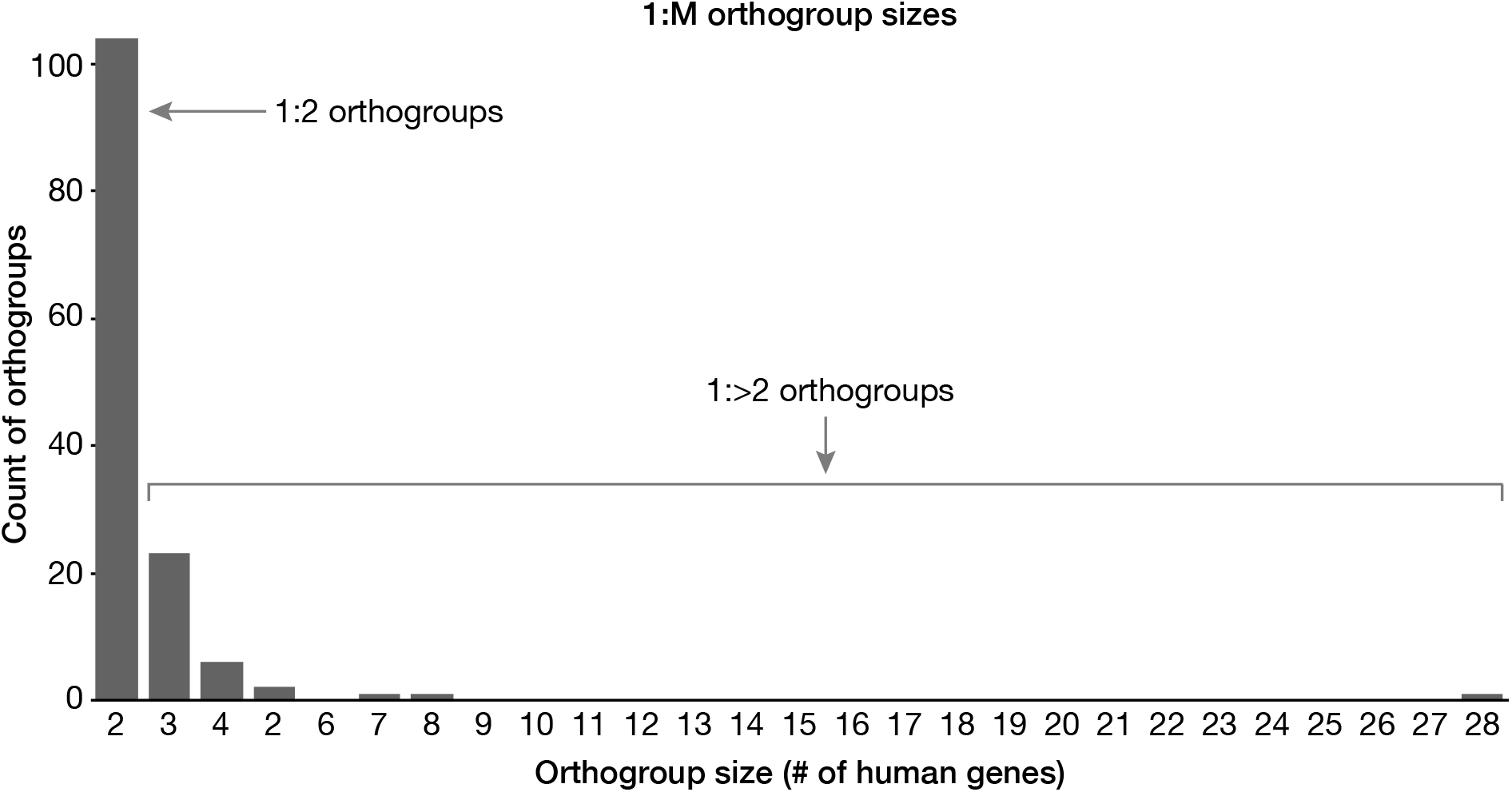
Count of orthogroups with the corresponding number of human gene members. The vast majority of 1 to multiple orthologs are of the 1:2 variety (1 yeast gene to 2 human co-orthologs) while the rest were classified as 1:>2 (1 yeast gene to >2 human co-orthologs). These two groups were considered separately during feature analysis.

**Figure S2.**
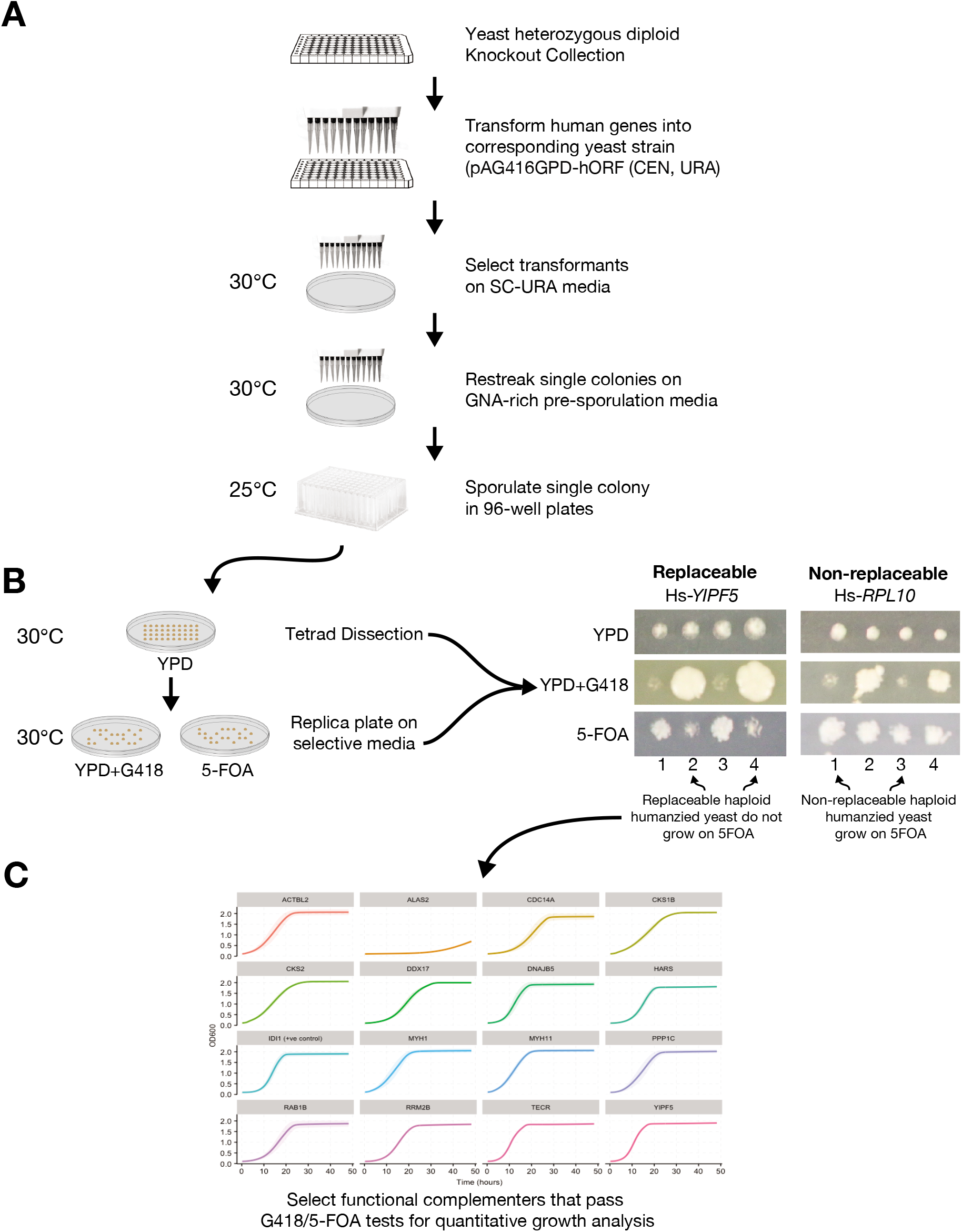
Detailed illustration of yeast gene replaceability, tetrad dissection and plasmid loss assays performed in yeast heterozygous diploid magic marker collection. **(A)** Yeast strains were grown in 96-well plates in selective medium (YPD + 200 *μ*g/ml G418). The matched orthologous human gene expression clones in 96-well plates were transformed into the appropriate yeast strains followed by selection on the appropriate dropout medium (SD-URA). The resulting transformants were spotted onto selective medium with appropriate markers to assay complementation. **(B)** In the case of the yeast Magic Marker heterozygous knockout collection, replacement was further verified by carrying out tetrad dissection followed by testing for plasmid dependency (selection on 5FOA). Representative tetrad dissection assays are shown (right). Hs-*YIPF5* functionally replaces its yeast counterpart. Dissected tetrads showed 2:2 segregation with tetrads 1 and 3 containing the wild-type yeast allele (surviving on 5FOA) while tetrads 2 and 4 contain the KanMX allele (yeast null allele) complemented by the human gene (surviving on YPD+G418). 5FOA and YPD+G418 are used for counter selecting the plasmid encoding the human gene and yeast null allele (KanMX selection) respectively. However, in the case of Hs-*RPL10* that fails to complement, yeast spore that grow on YPD+G418 also grow on 5FOA indicating plasmid loss. **(C)** Each confirmed humanized haploid yeast strain was assayed for growth defects to quantify the replaceability.

**Figure S3.**
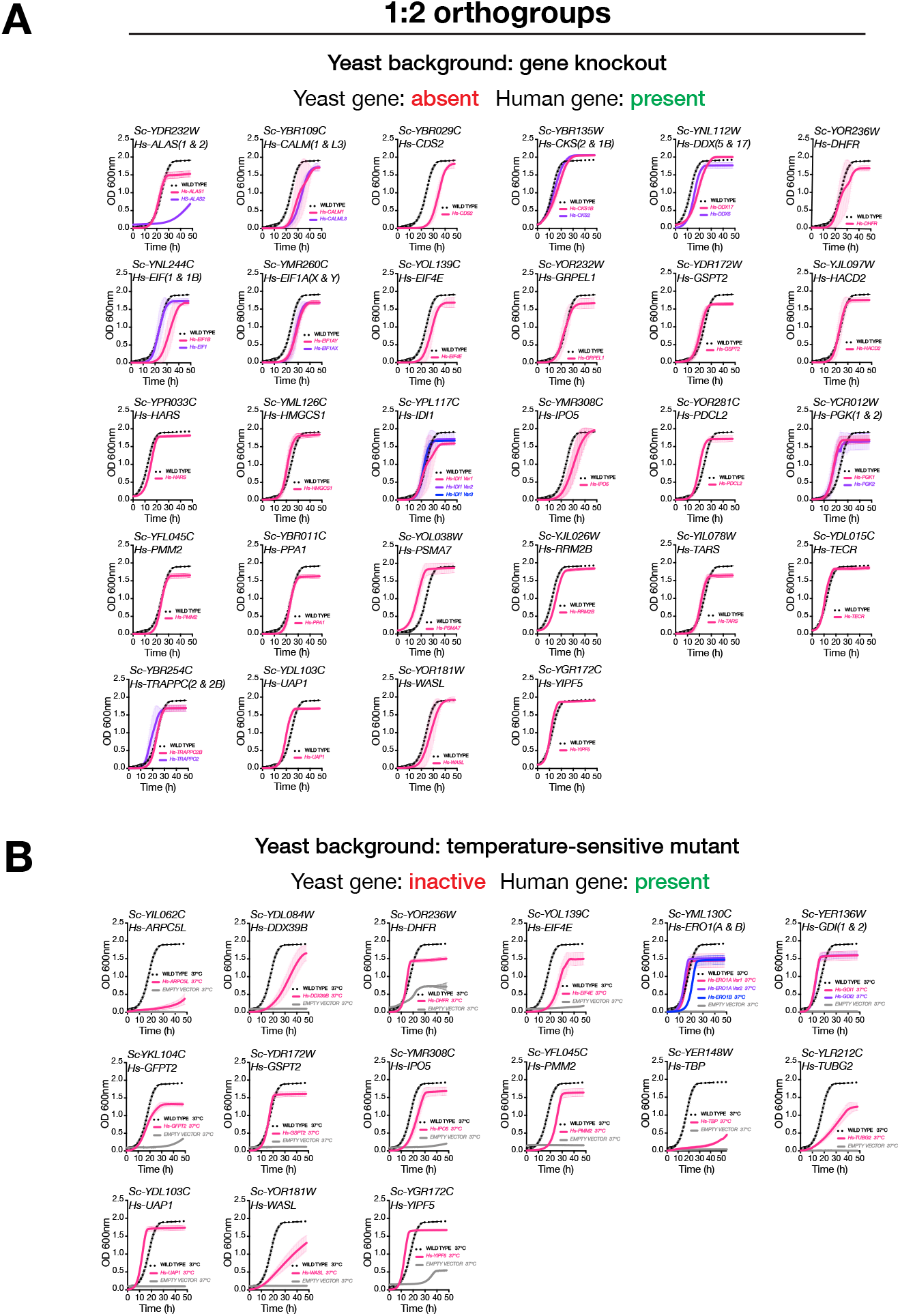
Quantitative growth assays of all humanized yeast strains belonging to 1:2 class. **(A)** Growth curves of humanized haploid yeast strains obtained after sporulation, tetrad dissection and followed by 5FOA screening of heterozygous diploid magic marker strains. The assays were performed in the presence of G418 (yeast gene-absent and human gene–present condition). Haploid yeast gene deletion strains carrying plasmids expressing functionally replacing human genes (colored red or purple solid-lines) generally exhibit comparable growth rates to the wild type parental yeast strain BY4741 (black dotted-lines) except in the case of Hs-*ALAS2* that shows reduced ability to replace the orthologous yeast gene compared to Hs-*AĨAS1.* Mean and standard deviation plotted from triplicate assays. **(B)** Growth curves of humanized haploid temperature sensitive yeast strains performed at restrictive temperature of 37°C (yeast gene – inactive and human gene – present condition). Temperature-sensitive haploid yeast strains carrying plasmids expressing functionally replacing human genes (colored red, purple or blue solid-lines) generally exhibit comparable growth rates to the wild type parental yeast strain BY4741 (black dotted-lines). The yeast strains harboring empty vector without a corresponding human gene, showing no or poor growth restrictive temperatures of 37°C (grey solid-lines), serve as controls. In total, 208 individual assays were performed (as two independent biological replicates). 45 human genes showed functionally replaced in either assay whereas 127 did not. 36 human gene complementation assays were non-informative (cases where the control experiments did not behave appropriately).

**Figure S4.**
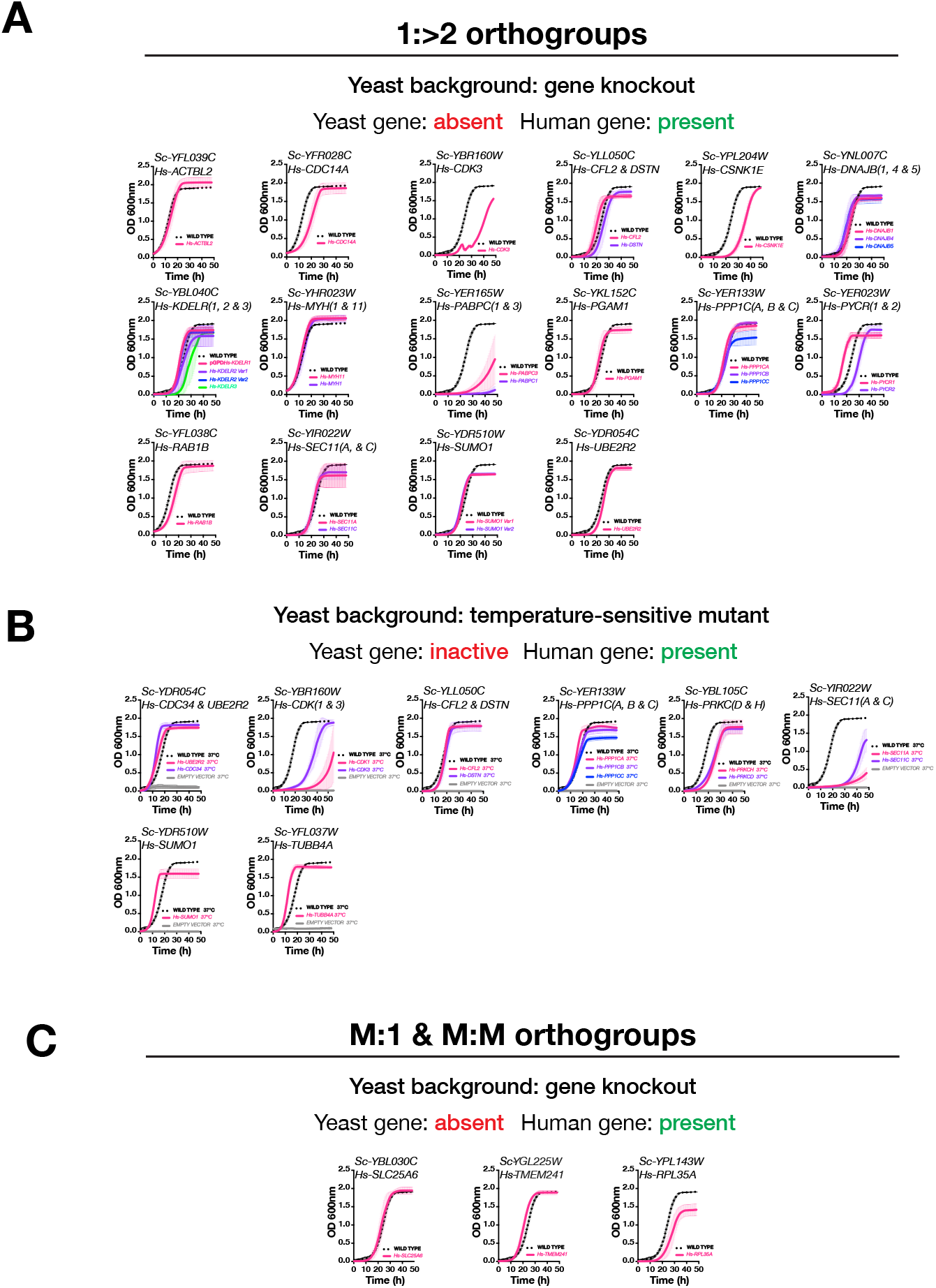
Quantitative growth assays of all humanized yeast strains belonging to 1:>2, M:M and M:1 orthogroups. **(A)** Growth curves of humanized haploid yeast strains (1:>2 class) obtained after sporulation and/or tetrad dissection followed by 5FOA screening of heterozygous diploid magic marker strains. The assays were performed in the presence of G418 (yeast gene-absent and human gene-present condition). Haploid yeast gene deletion strains carrying plasmids expressing functionally replacing human genes (colored red, purple and blue solid-lines) generally exhibit comparable growth rates to the wild type parental yeast strain BY4741 (black dotted-lines) except in the case of Hs-*PABPC1* and Hs-PABPC3 that shows reduced ability to replace the orthologous yeast gene. Mean and standard deviation plotted from triplicate assays. **(B)** Growth curves of humanized haploid temperature sensitive yeast strains (1:M class) performed at restrictive temperature of 37°C (yeast gene-inactive and human gene-present condition). Temperature-sensitive haploid yeast strains carrying plasmids expressing functionally replaced human genes (colored red, purple or blue solid-lines) generally exhibit growth rates comparable to its parental wild type yeast strain BY4741 (black dotted-lines) except in the cases of Hs-CDK3, *Hs-SEC11A* and *Hs-SEC11C* that showed reduced ability to functionally replace their orthologous yeast gene. Yeast strains harboring an empty vector without a corresponding human gene, showing no or poor growth at restrictive temperatures of 37°C (grey solid-lines), serve as controls. In total, 170 individual assays were performed (as two independent biological replicates). 32 human genes showed functional replaceability in either assay whereas 105 did not. 33 human gene replaceability assays were non-informative (cases when the control experiments did not behave appropriately). **(C)** Growth curves of humanized haploid yeast strains (M:1 and M:M class) obtained after sporulation and/or tetrad dissection and followed by 5FOA screening of heterozygous diploid magic marker strains. The assays were performed in the presence of G418 (yeast gene – absent and human gene – present condition). In total, 40 individual assays were performed (as two independent biological replicates). 3 human genes showed functional replaceability whereas 22 did not. 15 human gene replaceability assays were non-informative (cases when the control experiments did not behave appropriately). Mean and standard deviation plotted with N = 3.

**Figure S5.**
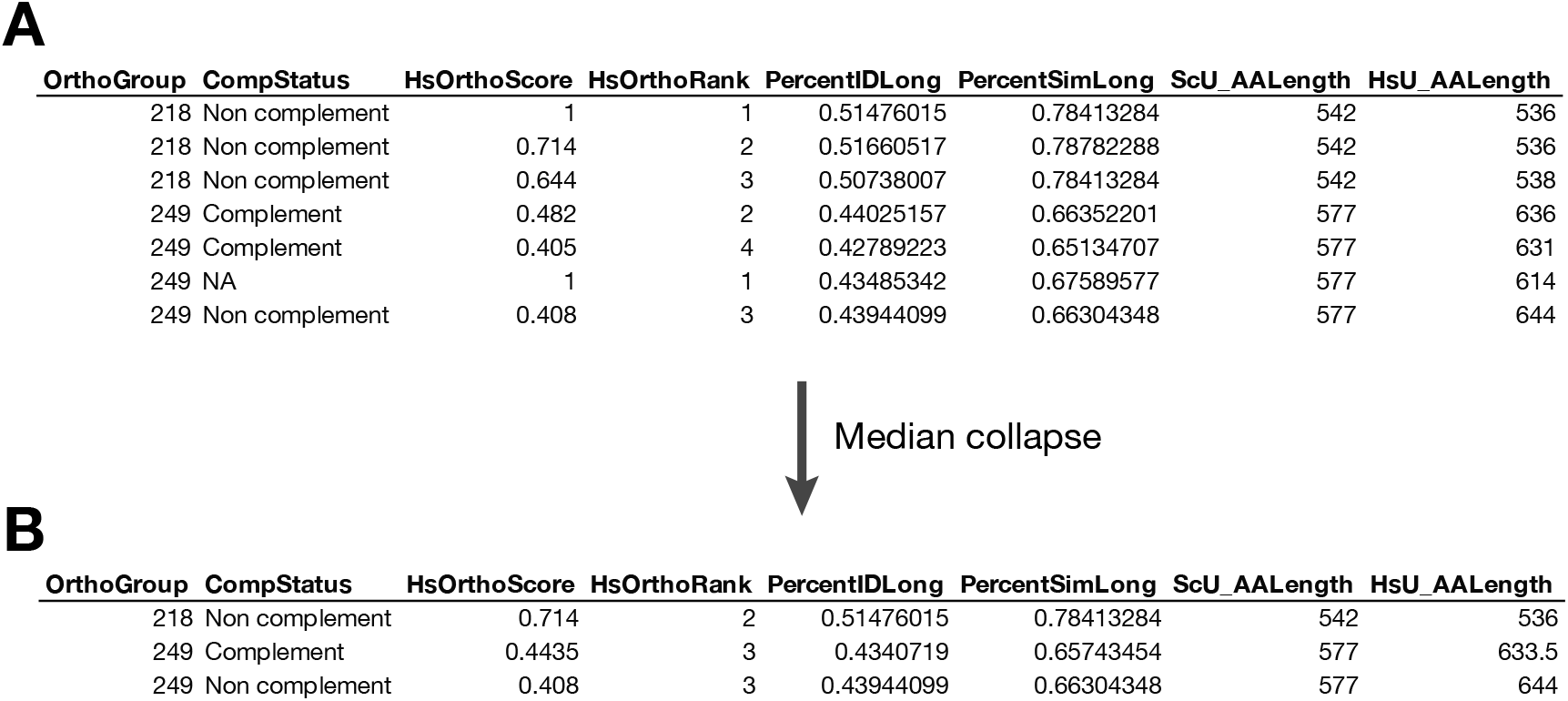
Demonstration of the ‘median-collapse’ feature table procedure. **(A)** An example subset of full ortholog data for two orthogroups in the 1:M ortholog class. **(B)** The same data as in **(A)** following median collapse. Orthogroup 218 has been collapsed to a single meta-ortholog pair with the status ‘Non complement’ as all human co-orthologs failed to complement. Group 249 has been collapsed to two meta-ortholog pairs, as there were co-orthologs that complemented and did not. Meta-ortholog feature values are the median of the full ortholog values with the same status.

